# Epidemiology and antimicrobial resistance characteristics of the ST131-*H30* subclone among extraintestinal *Escherichia coli* collected from US children

**DOI:** 10.1101/122416

**Authors:** Arianna Miles-Jay, Scott J. Weissman, Amanda L. Adler, Veronika Tchesnokova, Evgeni V. Sokurenko, Janet G. Baseman, Danielle M. Zerr

## Abstract

**Background:** *E. coli* ST131-*H30* is a globally important pathogen implicated in rising rates of multidrug resistance among *E. coli* causing extraintestinal infections. Previous studies have focused on adults, leaving the epidemiology of *H30* among children undefined.

**Methods:** We used clinical data and isolates from a case-control study of extended-spectrum cephalosporin-resistant *E. coli* conducted at four US children’s hospitals to estimate the burden and identify host correlates of infection with *H30. H30* isolates were identified using two-locus genotyping; host correlates were examined using log-binomial regression models stratified by extended-spectrum cephalosporin resistance status.

**Results:** A total of 339 extended-spectrum cephalosporin-resistant and 1008 extended-spectrum cephalosporin-susceptible *E. coli* isolates were available for analyses. The estimated period prevalence of *H30* was 5.3% among all extraintestinal *E. coli* isolates (95% confidence interval [CI] 4.6%-7.1%); *H30* made up 43.3% (81/187) of ESBL-producing isolates in this study. Host correlates of infection with *H30* differed by extended-spectrum cephalosporin resistance status: among resistant isolates, age ≤5 years was positively associated with *H30* infection (relative risk [RR] 1.83, 95% CI 1.19-2.83); among susceptible isolates, age ≤5 years was negatively associated with *H30* (RR 0.48, 95% CI 0.27-0.87), while presence of an underlying medical condition was positively associated (RR 4.49, 95% CI 2.43-8.31).

**Conclusions:** ST131-*H30* is less common among extraintestinal *E. coli* collected from children compared to reported estimates among adults, possibly reflecting infrequent fluoroquinolone use in pediatrics; however, it is similarly dominant among ESBL-producing isolates. The *H30* subclone appears to disproportionately affect young children relative to other extendedspectrum cephalosporin-resistant *E. coli*.

**Summary:** ST131-*H30* was responsible for 5.3% of all extraintestinal *E. coli* infections and 43.3% of ESBL-producing extraintestinal *E. coli* infections among US children. The clinical and demographic correlates of infection with ST131-*H30* differed between extended-spectrum cephalosporin-resistant and -sensitive isolates.

## INTRODUCTION

Extraintestinal *Escherichia coli*, a common cause of urinary tract and bloodstream infections across all ages, have displayed increasing rates of antimicrobial resistance over the past two decades.[1] This increase has been attributed to the emergence and rapid clonal expansion of *E. coli* Sequence Type (ST) 131, which has transformed the population structure of extraintestinal *E. coli* infections worldwide.[2–5] Molecular epidemiologic studies have shown that a subclone of ST131, termed *H30*, has driven the global dissemination of ST131.[6–9] The clonal structure of ST131-*H30* is tightly linked to antimicrobial resistance; the vast majority of *H30* isolates are fluoroquinolone resistant due to mutations in the *gyrA* and *parC* chromosomal genes (isolates known as *H30-R* or clade C), while nested subclones are additionally associated with the production of CTX-M-type extended-spectrum beta-lactamases (ESBLs) that confer resistance to extended-spectrum cephalosporins (Figure S1).[7,8,10–12]

Although *E. coli* ST131-*H30* (hereafter, *H30*) has been recognized as a clone of significant public health importance,[5,13] there is a lack of data about its epidemiology in children. Most studies that have included *H30* isolates from children have occurred over short time periods at single centers and have accumulated few *H30* isolates.[14–16] Among adults in the US, *H30* is estimated to comprise about 50% of ESBL-producing *E. coli* infections and 10%-20% of all extraintestinal *E. coli* infections, and has been linked to host factors including older age, healthcare contact, local or systemic compromise, and recent antibiotic use.[6,14–17] Associations with adverse outcomes such as persistent infections, new infections, sepsis, and hospitalization have also been reported in adult populations.[7,14,18] Understanding the epidemiology of *H30* in pediatric populations is important, as its dominance among multidrugresistant (MDR) extraintestinal *E. coli* makes it a likely culprit of many difficult-to-treat infections in children. Proper treatment of urinary tract infections – the most common type of infection caused by extraintestinal *E. coli* – is especially critical in pediatric populations, as young children are more prone to upper urinary tract infection with potential short- and long-term complications such as renal scarring and decreased renal function.[19,20]

We sought to address this knowledge gap using data from a multiyear, multicenter prospective case-control study of extraintestinal *E. coli* infections to quantify the burden and identify clinical and demographic correlates of infection with *H30* in a US pediatric population. In addition, we describe and compare the antimicrobial resistance characteristics of *H30* and non-*H30 E. coli* isolates.

## METHODS

### Patients and isolates

All isolates and clinical data came from a multicenter case-control study that prospectively collected isolates and is described in detail elsewhere.[21] In brief, between September 1, 2009 and September 30, 2013, four freestanding US children’s hospitals (referred to here as West, Midwest 1, Midwest 2, and East) used standard clinical microbiology techniques to identify and collect all extended-spectrum cephalosporin-resistant (ESC-R) *E. coli* collected from urine or other normally sterile sites during routine clinical care of both inpatient and outpatient children < 22 years of age. ESC-R isolates were defined as those non-susceptible to ceftriaxone, cefotaxime, ceftazidime, cefepime, or aztreonam. Patients could contribute multiple ESC-R isolates if the subsequent isolate was collected ≥ 15 days after the previous ESC-R isolate. For each resistant isolate, three consecutive *E. coli* isolates that were susceptible to the aforementioned agents, referred to here as extended-spectrum cephalosporin-susceptible (ESC-S) isolates, were collected without respect to any patient or microbiological characteristics beyond temporal proximity to the ESC-R isolates and prior enrollment in the study (patients could only contribute one ESC-S isolate). Demographic and clinical data were collected from the medical records; methods for categorizing underlying medical conditions, capturing antibiotic exposure, and characterizing the clinical significance of urine isolates (likely UTI vs. not) were described previously.[21,22] The Institutional Review Board at each hospital approved the study protocol.

### Laboratory methods

Methods for antibiotic susceptibility testing and typing of resistance phenotypes and determinants were described previously.[21] Briefly, ESC-R phenotypes (ESBL vs. AmpC) were characterized using a combination of disk diffusion and E-tests. Genetic determinants of extended-spectrum cephalosporin resistance were identified by PCR using primers for genes encoding common extended-spectrum cephalosporinases.[21] *H30* isolates were identified using the *fumC/fimH* genotyping scheme.[23] Isolates belonging to the *H30Rx* sublineage were identified by PCR detection of sublineage-specific single nucleotide polymorphisms.[7]

### Statistical analyses

#### Prevalence estimates

The period prevalence of *H30* was estimated by calculating a weighted average of the ESC-R and ESC-S stratum-specific prevalence estimates (details in Supplementary Methods).

#### Host correlates of infection

Only the first isolate from each unique individual was considered in the host factor analyses. Host factors were compared between patients with *H30* vs. non-*H30* isolates, stratified by ESC-R status and adjusting for study hospital where sample size allowed. The magnitude of the association between each predictor of interest and *H30* infection was then assessed using univariable and multivariable log-binomial regression models. For each predictor of interest, the relative risk (RR) and 95% confidence intervals (CIs) from three models are presented: 1) a univariable model that estimates the crude (unadjusted) total effect of the predictor of interest on the outcome; 2) a multivariable model that estimates the total effect of the predictor of interest on the outcome, adjusted for potential confounders; and 3) a multivariable model that estimates the direct effect of the predictor of interest on the outcome, adjusted for potential confounders as well as for potential mediators. All multivariable models adjusted for study hospital; additional potential confounders and mediators were selected according to the conceptual frameworks found in the supplementary material (Figures S3 & S4). Finally, we conducted *post-hoc* analyses of the interaction between age and underlying medical condition (details in Supplementary Methods).

#### Antimicrobial resistance characteristics

We examined co-resistance to commonly used antimicrobial agents in the first *E. coli* isolate collected per individual, stratifying by ESC-R and ESC-S status to maintain consistency with the sampling scheme of the parent study. *H30* isolates were additionally stratified into *H30Rx* and *H30-*non*-Rx* (Figure S1) and compared to non-*H30* isolates. Among ESC-R isolates, ESC-Rassociated resistance mechanisms and determinants were also identifed and compared. All analyses were conducted using R version 3.3.1 (R Core Team, 2016).

## RESULTS

### Isolates and prevalence estimates

A total of 339 ESC-R isolates from 278 patients and 1008 ESC-S isolates from 1008 patients were available for analyses (Figure S2). The estimated prevalence of H30 among all clinical E. coli isolates at all study hospitals was 5.3% (95% CI 4.6%-7.1%), while the hospital-specific prevalence ranged from 2.7% to 6.2% (Figure 1). The estimated overall prevalence of H30Rx was 0.87% (95% CI 0.70%-1.7%).

**Figure 1:**
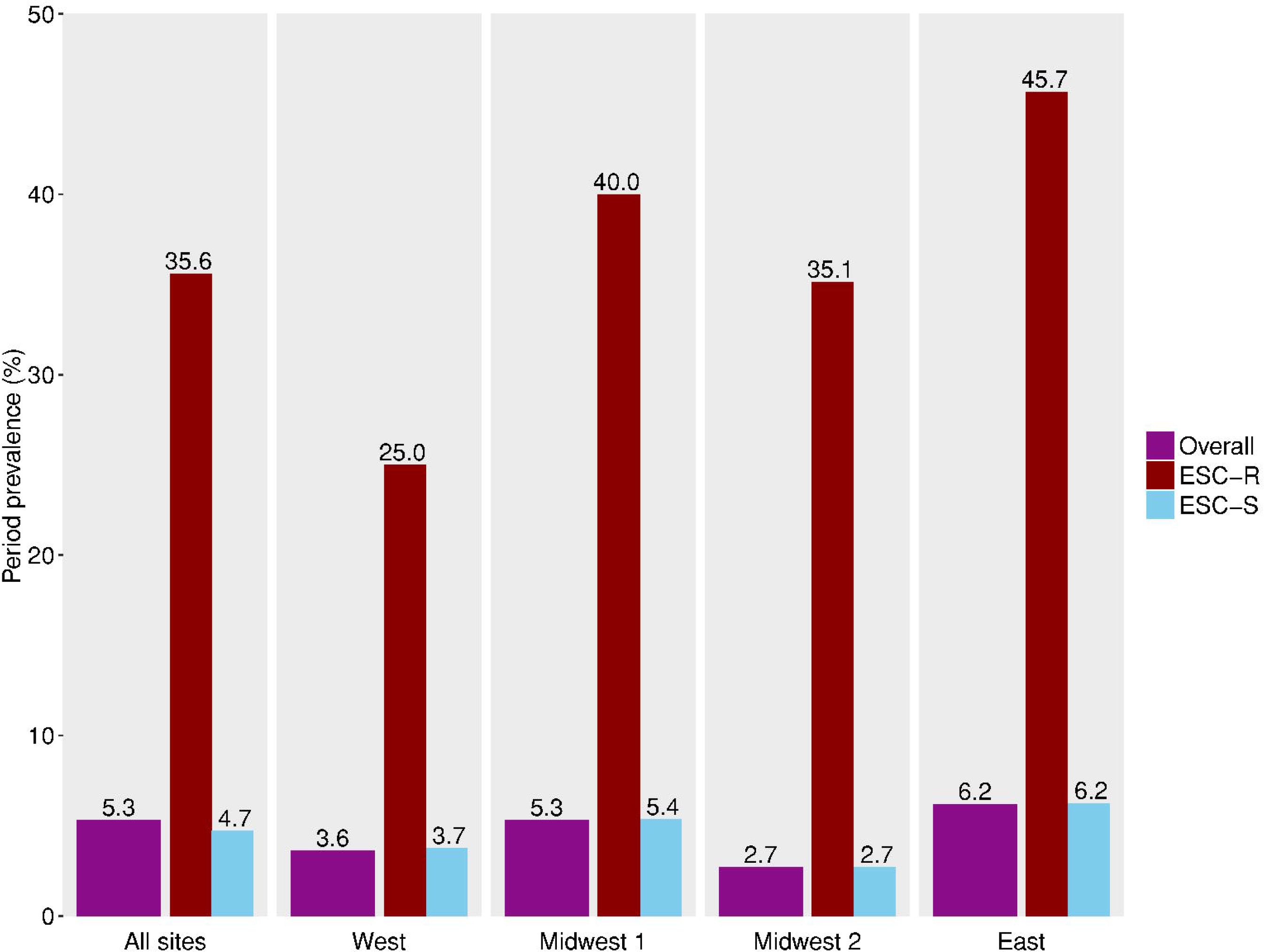
Estimated prevalence of ST131-*H30* among extraintestinal *E. coli* infections overall and by study hospital. ESC-R = extended-spectrum cephalosporin-resistant. ESC-S = extended-spectrum cephalosporin-susceptible. The raw numbers that generated these estimates can be found in Table S2.

### Host correlates of infection by ESC-R status

The first ESC-R isolate from each of the 278 patients with an ESC-R isolate collected during the study period was included in the host correlates analyses (Figure S2). Among these patients, patient age was associated with *H30* infection and further examined as a predictor of interest (Table 1). Our sample size precluded multilevel predictors, so age was categorized into ages 0-5 versus 6-21 years in regression models. After adjusting for potential confounders, age 0-5 was associated with an 83% increased risk of the infecting organism being *H30* (RR 1.83, 95%CI 1.19-2.83). There was no evidence that this association was mediated through factors related to underlying illness, or that underlying illness interacted with age (Table 3 and Table S3). When restricting the outcome to *H30Rx* infection only (vs. non-*H30* infection) and adjusting for potential confounders, the effect size was stronger (RR 2.25, 95%CI 1.33-3.80).

**Table 1:** Selected demographic and clinical characteristics of patients with *H30* and non-*H30* isolates, stratified by extended-spectrum cephalosporin resistance status

|  | ESC-R<br>n=278 |  |  | ESC-S<br>n=1008 |  |  |
| --- | --- | --- | --- | --- | --- | --- |
|  | *H30*<br>n=83 | non-*H30*<br>n=195 | p-value<sup>a</sup> | *H30*<br>n=47 | non-*H30*<br>n=961 | p-value<sup>a</sup> |
| **Age (years)** |  |  | 0.008* |  |  | <0.001* |
| 0-5 | 60 (72.3) | 98 (50.3) |  | 16 (34.0) | 504 (52.5) |  |
| 6-10 | 10 (12.1) | 40 (20.5) |  | 4 (8.5) | 190 (19.8) |  |
| 11-15 | 6 (7.2) | 31 (15.9) |  | 12 (25.5) | 126 (13.1) |  |
| 16-21 | 7 (8.4) | 26 (13.3) |  | 15 (31.9) | 141 (14.7) |  |
| **Sex** |  |  | 0.407 |  |  | 0.440 |
| Male | 18 (21.7) | 53 (27.2) |  | 8 (17.0) | 130 (13.5) |  |
| Female | 65 (78.3) | 142 (72.8) |  | 39 (83.0) | 831 (86.5) |  |
| **Ethnicity<sup>b</sup>** |  |  | 0.110 |  |  | 0.312 |
| Hispanic | 8 (10.0) | 36 (19.3) |  | 4 (8.9) | 135 (14.6) |  |
| Non-Hispanic | 72 (90.0) | 151 (80.7) |  | 41 (91.1) | 791 (85.4) |  |
| **Race<sup>b,c</sup>** |  |  | 0.087 |  |  | 0.314 |
| White/Caucasian | 39 (49.4) | 116 (62.0) |  | 29 (63.0) | 629 (68.2) |  |
| African-American | 12 (15.2) | 29 (15.5) |  | 12 (26.1) | 219 (23.7) |  |
| Asian | 22 (27.9) | 32 (17.1) |  | 2 (4.4) | 51 (5.5) |  |
| Native American | 4 (5.1) | 2 (1.1) |  | 1 (2.2) | 6 (0.7) |  |
| Pacific Islander | 2 (2.5) | 7 (3.7) |  | 1 (2.2) | 10 (1.1) |  |
| More than one race | 0 (--) | 1 (0.5) |  | 1 (2.2) | 8 (0.9) |  |
| **Site of culture** |  |  | 0.233 |  |  | 0.753 |
| Urine<sup>c,d</sup> | 78 (94.0) | 173 (88.7) |  | 45 (95.7) | 923 (96.1) |  |
| Blood | 2 (2.4) | 15 (7.7) |  | 2 (4.3) | 32 (3.3) |  |
| Other | 3 (3.6) | 7 (3.6) |  | 0 (0.0) | 6 (0.6) |  |
| **Type of acquisition<sup>b,e</sup>** |  |  | 0.832 |  |  | <0.001* |
| Community-associated | 28 (33.7) | 65 (33.3) |  | 14 (29.8) | 599 (62.7) |  |
| Healthcare-associated | 45 (54.2) | 103 (52.8) |  | 30 (63.8) | 297 (31.1) |  |
| Hospital-associated | 10 (12.0) | 27 (13.8) |  | 3 (6.4) | 60 (6.3) |  |
| **Hospitalized in past 6 months<sup>b</sup>** |  |  | 0.429 |  |  | 0.017* |
| Yes | 25 (30.1) | 69 (35.4) |  | 13 (27.7) | 143 (15.0) |  |
| No | 58 (69.9) | 126 (64.6) |  | 34 (72.3) | 813 (85.0) |  |
| **Underlying medical condition<sup>b</sup>** |  |  | 0.854 |  |  | <0.001* |
| Urologic<sup>f</sup> | 30 (36.1) | 75 (38.7) |  | 26 (55.3) | 185 (19.3) |  |
| Malignancy | 4 (4.8) | 13 (6.7) |  | 1 (2.1) | 26 (2.7) |  |
| Other condition | 16 (19.3) | 35 (18.0) |  | 6 (12.8) | 104 (10.8) |  |
| No condition | 33 (39.8) | 71 (36.6) |  | 14 (29.8) | 644 (67.2) |  |
| **Antibiotic use in the past 30 days<sup>b</sup>** |  |  | 0.548 |  |  | 0.006* |
| Yes | 34 (41.0) | 85 (43.6) |  | 16 (34.0) | 176 (18.4) |  |
| No | 49 (59.0) | 110 (56.4) |  | 31 (66.0) | 781 (81.6) |  |
| **History of transplantation<sup>b</sup>** |  |  | 0.108 |  |  | <0.001* |
| Yes | 3 (3.6) | 19 (9.7) |  | 5 (10.6) | 22 (2.3) |  |
| No | 80 (96.4) | 175 (90.2) |  | 42 (89.4) | 937 (97.7) |  |
| **Received immunosuppressants in last year<sup>b,g</sup>** |  |  | 0.100 |  |  | 0.071 |
| Yes | 9 (10.8) | 37 (19.1) |  | 6 (12.8) | 62 (6.5) |  |
| No | 74 (89.2) | 157 (80.9) |  | 41 (87.2) | 897 (93.5) |  |
| **Device type** |  |  | 0.157 |  |  | <0.001* |
| Central venous catheter | 7 (8.4) | 28 (14.4) |  | 3 (6.4) | 53 (5.5) |  |
| Foley catheter | 6 (7.2) | 5 (2.6) |  | 3 (6.4) | 11 (1.1) |  |
| Other device | 14 (16.9) | 26 (13.3) |  | 10 (21.3) | 55 (5.7) |  |
| No device | 56 (67.5) | 136 (69.7) |  | 31 (66.0) | 842 (87.6) |  |
| **Hospital** |  |  | 0.156 |  |  | 0.349 |
| West | 22 (26.5) | 78 (40.0) |  | 13 (27.7) | 341 (35.5) |  |
| East | 24 (28.9) | 51 (26.2) |  | 16 (34.0) | 284 (29.6) |  |
| Midwest 1 | 11 (13.3) | 23 (11.8) |  | 3 (6.4) | 108 (11.2) |  |
| Midwest 2 | 26 (31.3) | 43 (22.1) |  | 15 (31.9) | 228 (23.7) |  |

<sup>a</sup> P-values generated via Mantel-Haenzel chi-square tests (adjusting for study hospital) unless

**Table 2:** Selected antimicrobial resistance characteristics of *H30*Rx, *H30-non-Rx*, and non-*H30* isolates stratified by extendedspectrum cephalosporin resistance status

|  | ESC-R<br>n=278 |  |  |  |  | ESC-S<br>n=1008 |  |  |  |  |
| --- | --- | --- | --- | --- | --- | --- | --- | --- | --- | --- |
|  | *H30*<br>n = 83 |  | non-*H30*<br>n=195 | p-value vs. non-*H30*:<sup>a</sup> |  | *H30*<br>n = 47 |  | non-*H30*<br>n=961 | p-value vs. non-*H30*:<sup>a</sup> |  |
|  | Rx<br>n=64 | non-Rx<br>n=19 |  | Rx | non-Rx | Rx<br>n=5 | non-Rx<br>n=42 |  | Rx | non-Rx |
| **Co-resistance** |  |  |  |  |  |  |  |  |  |  |
| Ciprofloxacin | 62 (96.9) | 18 (94.7) | 76 (39.0) | <0.001* | <0.001* | 5 (100) | 36 (85.7) | 25 (2.6) | <0.001* | <0.001* |
| Gentamicin | 28 (43.8) | 6 (31.6) | 73 (37.4) | 0.453 | 0.798 | 0 (--) | 13 (31.0) | 34 (3.5) | 1.00 | <0.001* |
| TMP/SMX<sup>b</sup> | 43 (67.2) | 15 (78.9) | 121 (62.1) | 0.555 | 0.226 | 1 (20.0) | 26 (61.9) | 240 (25.0) | 1.00 | <0.001* |
| TMP/SMX & ciprofloxacin | 41 (64.1) | 15 (78.9) | 64 (32.8) | <0.001* | <0.001* | 1 (20.0) | 23 (54.8) | 15 (1.6) | 0.080 | <0.001* |
| All three | 19 (29.7) | 5 (26.3) | 36 (18.5) | 0.084 | 0.374 | 0 (--) | 8 (19.0) | 2 (0.2) | 1.00 | <0.001* |
| **ESC-R type** |  |  |  | <0.001* | 0.007* |  |  |  |  |  |
| ESBL only | 64 (100)<sup>c</sup> | 17 (89.5) | 102 (52.6) |  |  | -- | -- | -- | -- | -- |
| AmpC only | 0 (--) | 2 (5.4) | 88 (45.4) |  |  | -- | -- | -- | -- | -- |
| ESBL & AmpC | 0 (--) | 0 (--) | 4 (2.06) |  |  | -- | -- | -- | -- | -- |
| Undetermined | 0 (--) | 0 (--) | 1 (0.5) |  |  |  |  |  |  |  |
| **ESBL determinants<sup>d</sup>** | **n=64** | **n=17** | **n = 106** |  |  |  |  |  |  |  |
| CTX-M-15 | 60 (93.8)<sup>c</sup> | 3 (17.6) | 48 (45.3) | <0.001* | 0.060 | -- | -- | -- | -- | -- |
| CTX-M-14 | 0 (--) | 2 (11.8) | 44 (41.5) | <0.001* | 0.037* | -- | -- | -- | -- | -- |
| CTX-M-27 | 1 (1.6) | 10 (58.8) | 1 (0.9) | 1.000 | <0.001* | -- | -- | -- | -- | -- |
| CTX-M others | 0 (--) | 1 (5.3) | 7 (6.6) | 0.046* | 1.000 | -- | -- | -- | -- | -- |
| ESBL SHV | 0 (--) | 0 (--) | 3 (2.8) | 0.292 | 1.000 | -- | -- | -- | -- | -- |
| ESBL TEM | 0 (--) | 0 (--) | 0 (--) | -- | -- | -- | -- | -- | -- | -- |
| None identified | 3 (4.7) | 1 (5.3) | 4 (3.8) | 1.000 | 0.531 | -- | -- | -- | -- | -- |
| **AmpC determinants<sup>d</sup>** | **n=0** | **n=2** | **n=92** |  |  |  |  |  |  |  |
| CMY-2 | -- | 1 (50.0) | 79 (96.3) | -- | 0.277 | -- | -- | -- | -- | -- |
| DHA | -- | 0 (--) | 2 (2.2) | -- | 1.000 | -- | -- | -- | -- | -- |
| FOX | -- | 0 (--) | 2 (2.2) | -- | 1.000 | -- | -- | -- | -- | -- |
| None identified | -- | 1 (50.0) | 10 (10.9) | -- | 1.000 | -- | -- | -- | -- | -- |

^a^ P-values generated via Chi-square test; Fisher’s Exact test was used when expected frequencies were below 5. ^b^ TMP/SMX = trimethoprimsulfamethoxazole ^c^One of these isolates had both a CTX-M-15 gene identified as well as a KPC-3 carbapenemase gene, and was resistant to meropenem.^d^Total exceeds 100% as isolates could have more than one determinant identified. * p-value < 0.05

**Table 3:** Total and direct effect of selected factors on risk of *H30* infection vs. infection with other *E. coli* types using log-binomial regression models stratified by extended-spectrum cephalosporin resistance status.

|  | ESC-R |  |  | ESC-S |  |  |
| --- | --- | --- | --- | --- | --- | --- |
|  | Total effect<br>RR (95% CI) |  | Direct effect<br>RR (95% CI) | Total effect<br>RR (95% CI) |  | Direct effect<br>RR (95% CI) |
|  | Crude | Adjusted* | Adjusted* | Crude | Adjusted* | Adjusted* |
| **Age 0-5 years** | 1.98 (1.30-3.01)* | 1.83 (1.19-2.83)<sup>a†</sup> | 1.91 (1.24-2.96)<sup>b†</sup> | 0.48 (0.27-0.87)<sup>†</sup> | -- | 0.52 (0.29-0.94)<sup>c</sup> |
| **Antibiotics in last 30 days** | -- | -- | -- | 2.18 (1.22-3.91)<sup>†</sup> | 1.18 (0.64-2.20)<sup>d</sup> | -- |
| **Underlying medical condition** | -- | -- | -- | 4.46 (2.42-8.21)<sup>†</sup> | 4.49 (2.43-8.31)<sup>e†</sup> | 3.53 (1.74-7.17)<sup>f†</sup> |
| **Hospitalization in past 6 months** | -- | -- | -- | 2.08 (1.12-3.84)<sup>†</sup> | 1.22 (0.65-2.30)<sup>g</sup> | 1.01 (0.51-2.00)<sup>h</sup> |
| **Presence of indwelling device** | -- | -- | -- | 3.33 (1.87-5.92)<sup>†</sup> | 1.54 (0.78-3.04)<sup>i</sup> | 1.53 (0.77-3.01)<sup>c</sup> |

\*All models adjusted for study hospital.
<sup>a</sup> Additional covariates: Asian race (yes/no).

*All models adjusted for study hospital.
^a^ Additional covariates: Asian race (yes/no).
^b^ Additional covariates: Asian race (yes/no), underlying medical condition (yes/no), antibiotics in the last 30 days (yes/no), hospitalization in the past 6 months (yes/no).
^c^ Additional covariates: underlying medical condition (yes/no), antibiotics in the last 30 days (yes/no), hospitalization in the past 6 months (yes/no).
^d^ Additional covariates: age (0-5 or 6-21, hospitalization in the past 6 months (yes/no), underlying medical condition (yes/no), indwelling device (yes/no).
^e^ Additional covariates: age (0-5 or 6-21).
^f^ Additional covariates: age (0-5 or 6-21, hospitalization in the past 6 months (yes/no), antibiotics in the last 30 days (yes/no), indwelling device (yes/no).
^g^ Additional covariates: age (0-5 or 6-21), underlying medical condition (yes/no).
^h^ Additional covariates: age (0-5 or 6-21), underlying medical condition (yes/no), antibiotics in the last 30 days, indwelling device (yes/no).
^i^ Additional covariates: underlying medical condition (yes/no), hospitalization in the past 6 months (yes/no).
^†^Confidence interval does not include 1.

A total of 1008 patients had one ESC-S isolate collected during the study period. Among these patients, patient age and several factors associated with underlying illness were associated with *H30* infection (Table 1). Each of these variables was examined as a predictor of interest except for: (i) history of transplantation, due to small numbers, and (ii) type of infection acquisition, since previous hospitalization and underlying medical conditions were examined independently. Underlying medical condition and indwelling device categories were collapsed into any vs. none. Patient age ≤ 5 years was negatively associated with *H30* infection (RR 0.48, 95% CI 0.27-0.87). Of the variables related to underlying illness, after adjusting for potential confounders, only presence of an underlying medical condition (RR 4.49, 95%CI 2.43-8.31) remained as an independent predictor of *H30* infection; results were very similar when analyzing presence of an underlying urologic condition only (Table S4). When including potential mediators in the models, the magnitude of the associations between age ≤ 5 years and presence of an underlying medical condition with *H30* infection decreased, but the associations remained statistically significant (Table 3). Evidence of interaction between age and underlying medical condition was observed; when examining joint effects, underlying medical condition was only significantly associated with *H30* infection in combination with older age, and older age was only significantly associated with *H30* infection in combination with presence of an underlying medical condition (Table 4).

**Table 4:**
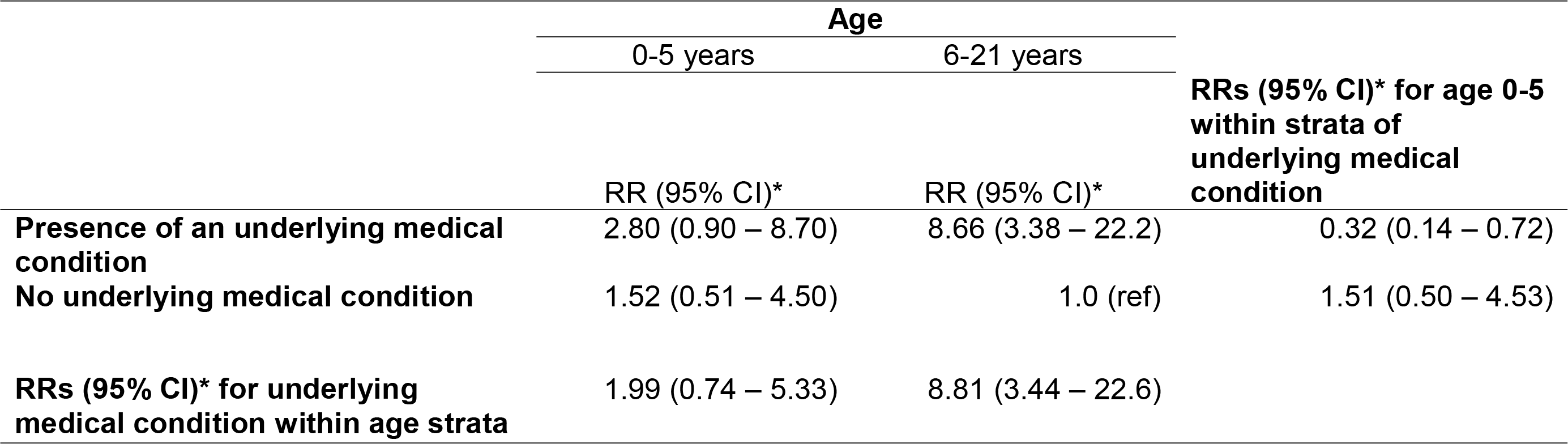
Analysis of interaction between age and underlying medical condition on the risk of *H30* infection vs. infection with other *E. coli* types using log-binomial regression models. Interaction Contrast Ratio (ICR) (95% CI) = −6.38 (−23.5 – −1.15). When interpreting the ICR, deviation from 0 indicates evidence of interaction on the additive scale (see Supplementary Methods). *RRs adjusted for study hospital.

|  | **Age** |  | **RRs (95% CI)\* for age 0-5 within strata of underlying medical condition** |
| --- | --- | --- | --- |
|  | 0-5 years | 6-21 years |  |
|  | RR (95% CI)* | RR (95% CI)* |  |
| **Presence of an underlying medical condition** | 2.80 (0.90 – 8.70) | 8.66 (3.38 – 22.2) | 0.32 (0.14 – 0.72) |
| **No underlying medical condition** | 1.52 (0.51 – 4.50) | 1.0 (ref) | 1.51 (0.50 – 4.53) |
| **RRs (95% CI)\* for underlying medical condition within age strata** | 1.99 (0.74 – 5.33) | 8.81 (3.44 – 22.6) |  |
| Interaction Contrast Ratio (ICR) (95% CI) = -6.38 (-23.5 – -1.15). When interpreting the ICR, deviation from 0 |  |  |  |

Since patient age was important in the analyses of both ESC-R and ESC-S isolates, we also visually inspected the distributional differences of age measured continuously. While the non-*H30* age distributions are very similar, the *H30* age distributions display marked differences between ESC-R and ESC-S isolates (Figure 2).

**Figure 2:**
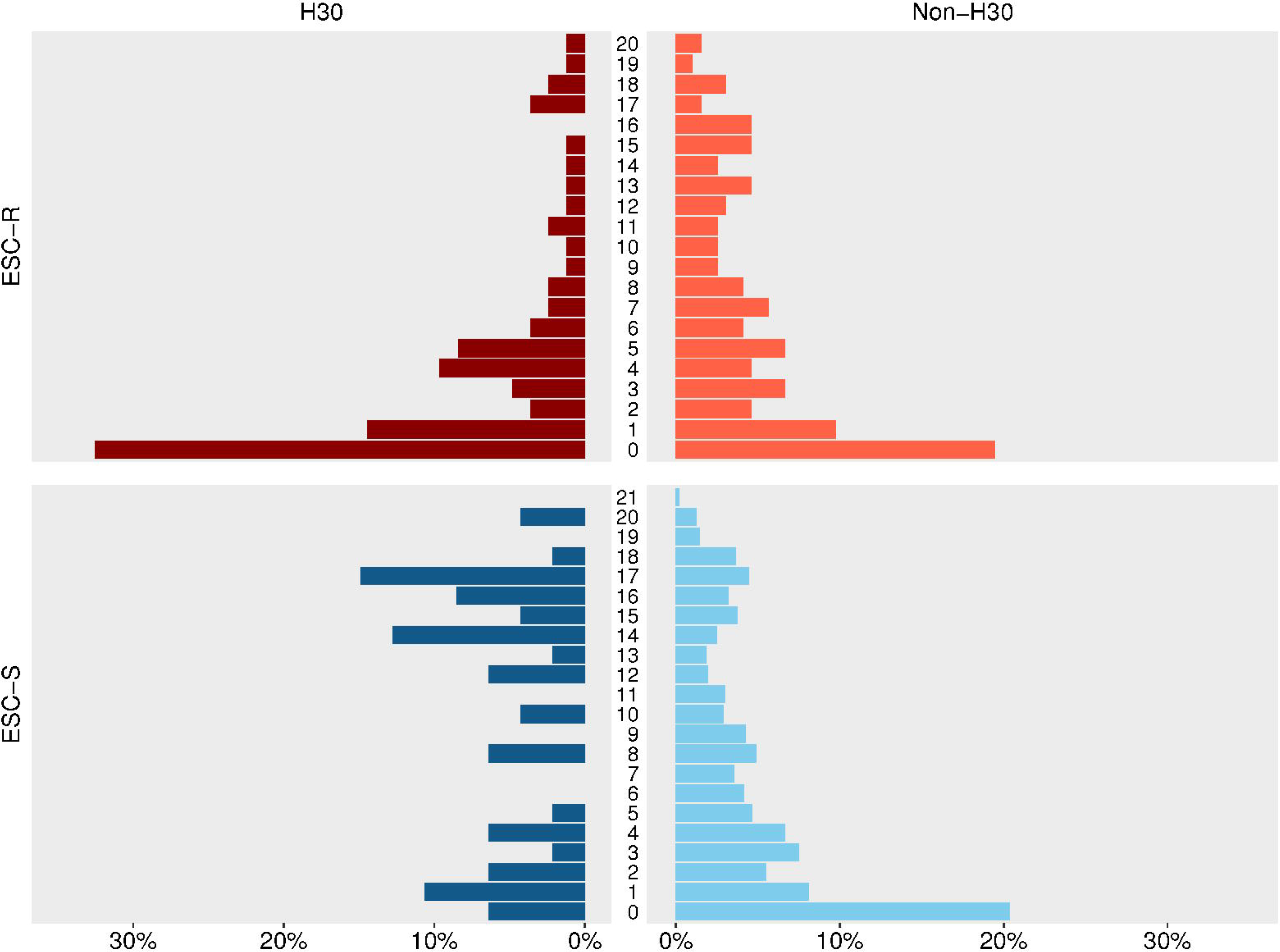
Distributions of age (in years) by ST131-*H30* and non-ST131-*H30* status and extended-spectrum cephalosporin resistance status. ESC-R = extended-spectrum cephalosporin-resistant. ESC-S = extended-spectrum cephalosporin-susceptible.

### Antimicrobial resistance characteristics by ESC-R and *H30Rx* status

A total of 278 ESC-R isolates were examined (the first isolate collected per individual). Among these isolates, nearly all *H30Rx* and *H30-*non*-Rx* isolates were non-susceptible to fluoroquinolones, compared to less than half of non-*H30* isolates (Table 2). Similarly, all ESC-R *H30Rx* and the vast majority of *H30-*non-*Rx* isolates were ESBL-producing, while non-*H30* isolates were more evenly split between ESBL producers and AmpC producers. *H30* was the most common subclone identified among the ESC-R isolates in the study (Table S1); it made up 29.9% (83/278) of ESC-R isolates, and when restricting to ESBL-producing isolates only, it made up 43.3% (81/187) of the total. The vast majority of ESBL-producing *H30Rx* isolates had a CTX-M-15 beta-lactamase, while ESBL-producing *H30-*non*-Rx* isolates were dominated by the CTX-M-27 beta-lactamase; ESBL-producing non-*H30* isolates were more evenly split between CTX-M-15 and CTX-M-14 beta-lactamases (Table 2). Systematic differences in the types of ESC-R resistance determinants by study hospital or year were not observed (Figure S5).

Among the 1008 ESC-S isolates examined, fluoroquinolone non-susceptibility was dominant among *H30* isolates, while only a small fraction of non-*H30* ESC-S isolates were nonsusceptible to fluoroquinolones (Table 2).

## DISCUSSION

We utilized a multiyear, multicenter case-control study of extraintestinal *E. coli* infections in children’s hospitals to address a critical knowledge gap about the epidemiology of the globally important ST131-*H30* subclone among US children. Our results can be summarized into three main findings. First, the estimated prevalence of *H30* among pediatric extraintestinal *E. coli* isolates of 5.3% was lower than the 10-20% that has been observed in US adults.[6,14,15] However, *H30* was nearly as dominant among ESBL-producing isolates in children (43.3%) as has been reported in adults (about 50%).[16,17] Second, patient age was associated with infection due to *H30*, and the nature of this association contrasted sharply between ESC-R and ESC-S infections. Among ESC-R infections, *H30* was associated with young age (≤5 years), while among ESC-S infections, *H30* was associated with older age (6-21 years), as well as with the presence of an underlying medical condition. Third, the antimicrobial resistance characteristics of *H30* and *H30Rx* collected from children were consistent with what has been previously reported.[12,16–18,24] ESC-R *H30* isolates were almost always fluoroquinolone-resistant and ESBL-producing, and ESBL-producing *H30Rx* isolates were associated with the CTX-M-15 beta-lactamase, while ESBL-producing *H30-*non*-Rx* isolates were associated with the CTX-M-27 beta-lactamase.

Other studies have suggested that *H30* is less prevalent among children than adults; however, very few pediatric isolates were included in these studies.[15,16] Interestingly, we observed that *H30* was nearly as dominant among ESBL-producing *E. coli* infections in children as has been reported in adults.[16,17] These findings are consistent with a recent study from a pediatric setting conducted in the Midwestern US.[25] However, in the context of all clinical extraintestinal *E. coli* infections, ESBL-producing organisms are still relatively rare in both adults and children. The bulk of the *H30* isolates circulating in the population are non-ESBL-producing but fluoroquinolone-resistant, and these isolates were much less common in our study than has been observed in adult populations.[15,16] This observation may be explained by differential antibiotic use in these populations. Fluoroquinolones are infrequently prescribed to children due to concerns about toxicity;[26] in our study, about 5% of patients received fluoroquinolones in the year before collection of their first isolate, while 46% of patients received any antibiotic in that same time period (Figure S6). Lower rates of fluoroquinolone use likely translate to less selective pressure on fluoroquinolone-resistant organisms such as *H30*. Interestingly, a recent study conducted in adults in Australia and New Zealand, a population that also has low rates of fluoroquinolone use, reported an overall prevalence of *H30* of 3.5%, but a prevalence of *H30* among ESC-R *E. coli* of 39%, which is similar to our findings.[27]

The association we identified between *H30* and young age among ESC-R isolates is consistent with the findings of a recent longitudinal study showing that among children, the prevalence of ESBL-producing *Enterobacteriaceae* was highest and increasing most rapidly in children aged 1-5.[28] Why *H30/H30Rx* is more frequently found among young children with ESC-R infections compared to older children with ESC-R infections, as well as where young children are acquiring this pathogen, deserves further investigation. Previous studies have portrayed *H30* as an opportunistic pathogen that favors compromised hosts including the elderly,[14] and young children’s developing immune systems could be associated with *H30* infection. Maternal infection or colonization may also play a role; a recent study found *H30* colonization during the first several years of life of healthy twins was associated with the mother also being colonized, however, none of these *H30* isolates were ESBL-producing.[29] Finally, while transmission of *H30* between children within healthcare facilities has not been documented, there are reports of transmission of, and persistent colonization with, *H30/H30Rx* among healthy children within daycares and households.[29–32] Future studies might focus on systematic sampling in the community setting in order to better elucidate the reservoirs and transmission dynamics of *H30/H30Rx* among young children.

The association we observed between ESC-S *H30* infections and older children is not consistent with the limited existing data.[15,33]. Our *post-hoc* interaction analyses suggest that age and underlying illness interact, with the strongest risk of an infection being *H30* observed in older children with underlying medical conditions. We hypothesize that these observed associations could be driven by different selective pressures in older, less healthy children: specifically, fluoroquinolones are likely prescribed more frequently to older children than younger children due to less concern about toxicity. This prescribing pattern was borne out in our data; the median age was 12.6 years among patients that received fluoroquinolones in the year prior to their infection, whereas the median age among those that received any antibiotic was 6 years (Figure S6). A more refined examination of the role of antibiotic exposure, specifically focusing on fluoroquinolones, is warranted.

Notably, previous studies conducted in adult populations have described *H30* as being associated with healthcare contact and compromised hosts,[14,15] however, we found those associations only among ESC-S *H30* infections. The fact that we observed these patterns among ESC-S isolates is not surprising; compromised hosts and healthcare contact are consistently associated with antimicrobial resistant infections,[34] and as is shown in Table 2, *H30* isolates are more antimicrobial-resistant than other ESC-S isolates. However, we observed that when compared to other ESC-R organisms, there is no evidence of an association between *H30* and underlying illness. This observation raises the question of whether some host correlates observed in previous studies are specific to the *H30* subclone, or just reflect risk factors for MDR extraintestinal *E. coli* in general. Future studies should consider comparing *H30* to other MDR *E. coli* where possible.

A number of limitations need to be considered in the interpretation of these data. First, because of the case-control design of the parent study, the prevalence of *H30* and *H30*Rx among clinical *E. coli* isolates could not be calculated directly. However, we believe the assumptions employed in our prevalence estimates are reasonable, and that these data provide the best estimate of the prevalence of *H30* in children to date. The design of the parent study was also a strength, as it allowed us to enrich the collection with the less common MDR isolates and examine risk factors for infection with *H30* among those with ESC-R *E. coli* isolates specifically. Second, because this study was an exploratory investigation of an existing dataset, all findings should be interpreted cautiously; there could be residual confounding due to unmeasured or incompletely measured variables, spurious associations identified due to multiple testing, or missed associations due to lack of power. To mitigate this, we attempted to make thoughtful model building decisions and interpretations by using conceptual models rather than taking a purely data-driven approach. Third, the isolates did not undergo multilocus sequence-typing (MLST) or other molecular characterization relevant to *H30* such as typing of the *gyrA* and *parC* alleles. However, the *H30* isolates in this study have since undergone whole genome sequencing, and *in silico* MLST analyses have confirmed that isolates classified as *H30* are ST131 (data not shown). Finally, although this was a multicenter study, our data were collected from freestanding children’s hospitals between 2009 and 2013, so the results may not be generalizable to other settings, and epidemiologic patterns may have shifted during the subsequent several years. Despite these limitations, this study significantly improves our understanding of the impact of *H30* in children, and is one of the most robust examinations of the clinical burden of, and risk factors for, *H30* infections to date.

### Conclusion

Although *E. coli* ST131-*H30* is not as prevalent among children as has been reported in adults, perhaps as a result of low rates of fluoroquinolone use in pediatrics, this clone is dominant among ESC-R extraintestinal *E. coli* infections in children. In particular, ESBL-producing *H30*, dominated by the *H30Rx* subclone, disproportionately affect young children relative to other ESC-R *E. coli*, even when accounting for other underlying host factors. More densely sampled studies are needed to elucidate the reservoirs and transmission dynamics of this difficult-to-treat pathogen in a pediatric population.

## Acknowledgements

The authors thank Carey-Ann Burnham, Alexis Elward, Jason Newland, Rangaraj Selvarangan, Kaede Sullivan, Theoklis Zaoutis, and Xuan Qin for their provision of bacterial isolates and associated clinical data. Additionally, they thank Jeff Myers and Huxley Smart for their assistance with molecular typing of isolates.

